# A screen for gene paralogies delineating evolutionary branching order of early Metazoa

**DOI:** 10.1101/704551

**Authors:** Albert Erives, Bernd Fritzsch

## Abstract

The evolutionary diversification of animals is one of Earth’s greatest triumphs, yet its origins are still shrouded in mystery. Animals, the monophyletic clade known as Metazoa, evolved wildly divergent multicellular life strategies featuring ciliated sensory epithelia. In many lineages epithelial sensoria became coupled to increasingly complex nervous systems. Currently, different phylogenetic analyses of single-copy genes support mutually-exclusive possibilities that either Porifera or Ctenophora is sister to all other animals. Resolving this dilemma would advance the ecological and evolutionary understanding of the first animals and the evolution of nervous systems. Here we describe a comparative phylogenetic approach based on gene duplications. We computationally identify and analyze gene families with early metazoan duplications using an approach that mitigates apparent gene loss resulting from the miscalling of paralogs. In the transmembrane channel-like (TMC) family of mechano-transducing channels, we find ancient duplications that define separate clades for Eumetazoa (Placozoa + Cnidaria + Bilateria) versus Ctenophora, and one duplication that is shared only by Eumetazoa and Porifera. In the MLX/MLXIP family of bHLH-ZIP regulators of metabolism, we find that all major lineages from Eumetazoa and Porifera (sponges) share a duplication, absent in Ctenophora. These results suggest a new avenue for deducing deep phylogeny by choosing rather than avoiding ancient gene paralogies.

## INTRODUCTION

The branching order of the major metazoan lineages has received much attention due to its importance in understanding how animals and their sensory and nervous systems evolved (Jekely *et al*. 2015). To date, most phylogenetic analyses have used single-copy orthologs, with different genes and approaches finding support for either Ctenophora (Ryan *et al*. 2013; Moroz *et al*. 2014; Borowiec *et al*. 2015; Whelan *et al*. 2015; Shen *et al*. 2017) or Porifera (Pisani *et al*. 2015; Feuda *et al*. 2017; Simion *et al*. 2017) as the sister taxon of all other animals. In contrast to sequence-based phylogenies, comparative analysis of single-cell transcriptomes from different lineages and cell types is consistent with an independent origin of neuron-like cells in ctenophores (Sebe-Pedros *et al*. 2018). Another recent study that does not rely directly on sequence analysis provides evidence that the sponge choanocyte does not correspond to the sponge cell type most similar transcriptomically to the choanoflagellate cell type (Sogabe *et al*. 2019). Thus to date, few studies (Sebe-Pedros *et al*. 2018; Sogabe *et al*. 2019) have addressed the question of early animal branching without using single-copy genes for phylogenetic analysis.

Single-copy genes are preferred for phylogenetic inference of lineage branching order for several reasons (Fitch and Margoliash 1967; Woese and Fox 1977). For example, single-copy genes more closely approximate clock-like divergence (Zuckerkandl and Pauling 1965; Fitch and Margoliash 1967; Woese and Fox 1977; Pett *et al*. 2019) compared to duplicated genes, which frequently experience neofunctionalization and/or uneven subfunctionalization and evolutionary rate asymmetries (Walsh 1995; Lynch *et al*. 2001; Holland *et al*. 2017). An organismal tree of animals can be constructed from a single-copy gene, or from a set of concatenated single-copy genes, provided orthologous outgroup genes are included in the analysis as an aid for rooting.

Likewise, single-copy genes (strict orthologs across all lineages) offer a practical advantage in eliminating the ambiguity associated with gene duplications (paralogs). Some paralogs represent recent lineage-specific duplications while others stem from deeper duplications contributing to a larger gene super-family. Nonetheless, a gene tree of a pair of genes produced by a duplication in the stem-metazoan lineage can depict lineage-branching order doubly so, once in each paralog’s subtree. Furthermore, a gene tree of paralogs offers a unique advantage unavailable in single-copy gene trees: a tree of paralogs captures the duplication itself and unites lineages sharing the duplication relative to outgroup lineages with a single gene.

A majority of gene orthologs are part of larger super-families (Ohno 1970; Taylor and Raes 2004) as has been well documented for the Hox gene family (Holland *et al*. 2017). Thus, many “single-copy” genes are only ostensibly so because they can be evaluated separately from their ancient paralogs; and because in principle single-copy genes diverged lineally from a single ancestral gene present in the latest common ancestor (LCA) of a taxonomic clade.

However, the choice of homologs is a poorly examined aspect of modern phylogenetic analysis even though various ortholog-calling errors associated with ancient paralogy have been noted (Noutahi *et al*. 2016). Here we identify candidate gene paralogies established prior to the evolution of Eumetazoa (Bilateria + Cnidaria + Placozoa) for the purpose of determining early animal branching order. The majority of these paralogies predate the metazoan LCA and/or experienced apparent gene losses in either candidate first animal sister lineage (Porifera or Ctenophora) and are not informative (see Table S1). We also find a smaller number of paralogies that possibly support a proposed clade of “Benthozoa” (Porifera + Eumetazoa) while we have found none that support the traditional grouping of Ctenophora with Eumetazoa. The benthozoic hypothesis is premised on the latest common ancestor (LCA) of Choanozoa (Choanoflagellatea + Metazoa) being holopelagic, and the LCA of Benthozoa having evolved a biphasic pelagic larva and a benthic adult form. Other alternative life cycle scenarios have been proposed that correspond to different branching patterns (Nielsen 2008; Nielsen 2013; Jekely *et al*. 2015) some of which are based on fossil interpretation (Zhao *et al*. 2019).

Among the most intriguing of our findings are the recently identified family of multimeric transmembrane mechanosensitive channel proteins, the transmembrane channel-like (TMC) proteins (Keresztes *et al*. 2003; Ballesteros *et al*. 2018), in which family we find definitive independent duplications in Eumetazoa (*Tmc<u>48756</u>* → *Tmc<u>487</u>* + *Tmc<u>56</u>*), Ctenophora (*Tmc<u>48756</u>* → *Tmc-α*, *Tmc-β*, *Tmc-γ*, and *Tmc-δ*), and possibly Benthozoa (*Tmc<u>12348756</u>* → *Tmc<u>48756</u>* + a neofunctionalized *Tmc<u>123</u>* clade). In summary, our identification and analysis of genes duplicating and diversifying in the stem-benthozoic lineage will help to outline the extent of a shared biology for Benthozoa and the nature of independent evolutionary neuralization in Eumetazoa and Ctenophora.

## Material & Methods

### Comparative genomic orthology counting and screening

For the data depicted in Fig. 3, we used BioMart query tool (Durinck *et al*. 2005; Haider *et al*. 2009; Smedley *et al*. 2009) and the EnsemblCompara orthology calls (Vilella *et al*. 2009) for MetazoaEnsembl Genomes Release 41. We also used these same tools to identify 2146 unique protein-coding genes in the placozoan *Trichoplax adhaerens*, which share the following properties: (1) these genes all can be grouped into a smaller number of paralogy groups; (2) these genes all have homologs in the cnidarian *Nematostella vectensis*; (3) the cnidarian homologs can also all be grouped into a smaller number of paralogy groups; (4) these genes all have homologs in the molluscan genome of *Lottia gigantea*, representing Lophotrochozoa, as well as in *Drosophila melanogaster*, representing Ecdysozoa; and (5) these genes all have homologs in the sponge *Amphimedon queenslandica* (a sponge in class Demospongiae). To ensure that our results would not be skewed by errors in gene annotation and curation, we focused on genes for which we could identify homologs in other cnidarians (the anthozoan *Stylophora pistillata* and the hydrozoan *Hydra vulgaris*), another sponge (the homoscleromorphan sponge *Oscarella carmela*), and throughout Bilateria. We then constructed phylogenetic trees for several different gene families from this list.

### Sequence alignment

We identified and curated sequences only from representative taxa with whole-genome sequence assemblies. We obtained initial (pre-full-length curation) sequences from NCBI’s non-redundant protein database using the BLAST query tool with taxonomic specification (Altschul *et al*. 1990), and/or from the Ensembl Release 97 and Metazoa Ensembl Release 45 databases using the ComparaEnsembl orthology calls (Vilella *et al*. 2009). For additional ctenophore sequences from the *Pleurobrachia* and *Beröe* genomes, we queried the Neurobase transcriptome databases using BLAST (Moroz *et al*. 2014). For additional sequences and transcripts from cnidarian and sponge genomes we also queried the Compagen databases (Hemmrich and Bosch 2008). In addition to BLASTP queries of protein or translated transcriptome databases, we also used the TBLASTN tool to search translated genomic sequences using a protein sequence. We did this for two reasons. First, we used TBLASTN to verify that certain gene absences were not simply due to a failure to annotate a gene. Second, we used TBLASTN on several occasions when curating missing exons. We identified several genes from genome assemblies that were initially predicted by computational annotation and for which we then hand-curated for this study, typically to identify missing terminal exons. These are indicated in the tree figures and in the FASTA headers (Supplementary files) by the accession numbers with “CUR” appended. Alignment of protein-coding sequence was conducted using MUSCLE alignment option in MEGA7 (Kumar *et al*. 2016). The multiple sequence alignment (MSA) was conducted using default parameters that were adjusted as follows. The gap existence parameter was changed to −1.6 and the gap extension parameter was changed to −0.01. Excessively long protein sequences were trimmed at the N- and C-termini so that they began and ended on either side of the ten transmembrane domains. Lengthy, fast-evolving, loop segments and/or repetitive amino acid sequences occurring in between transmembrane domains were trimmed. Supplementary Files for curated data sets are provided as explained under “Data availability”.

### Bayesian Inference

To conduct metropolis-coupled-MCMC Bayesian phylogenetic we used the MrBayes (version 3.2) software (Huelsenbeck and Ronquist 2001; Ronquist and Huelsenbeck 2003; Ronquist *et al*. 2012). All runs used two heated chains (“nchains = 2”, “temperature = 0.08”) that were run for 1.2 M generations with a burn-in fraction of 20%. Initial runs for all gene families sampled all substitution models, but we always found 100% percent posterior probability assigned to the WAG substitution model (Whelan and Goldman 2001). Subsequently all finishing runs used the WAG model with invariant-gamma rates modeling. Double precision computing was enabled using BEAGLE toolkit (Ayres *et al*. 2012; Ronquist *et al*. 2012). All trees were computed multiple times during the process of sequence curation and annotation. The final *cornichon*/*cornichon-related* gene family tree of Figure 4B (ver. 21) finished with 0.009 average standard deviation of split frequencies from two heated chains. The final *Tmc* gene family tree of Figure 5 (ver. 69) finished with 0.005 average standard deviation of split frequencies from the two heated chains. The final *MLX*/*MLXIP* gene family tree of Figure 6 (ver. 20) finished with 0.007 average standard deviation of split frequencies from the two heated chains. Tree graphics were rendered with FigTree version 1.4.4 and annotated with Adobe Photoshop tools.

### Data availability

All data files, including sequence files (*.fas), multiple sequence alignment files (*.masx and *.nexus), and curation documents (Curated_*_Genes.docx), are provided with the Supporting Information as a zipped archive. Four files are provided for each of the three gene families shown for a total of 12 files.

## RESULTS

### Gene duplications from the early metazoan radiation

Given the preponderance and constancy of gene duplication (and gene loss) throughout evolution, one should in principle be able to find a gene duplication shared by all major animal lineages except the one true sister animal lineage. Therefore, ever since the elucidation of the first non-bilaterian animal genomes, mainly those from Cnidaria (Putnam *et al*. 2007), Placozoa (Srivastava *et al*. 2008), Porifera (Srivastava *et al*. 2010), and Ctenophora (Ryan *et al*. 2013; Moroz *et al*. 2014), we have sought to identify gene duplications that definitively order early metazoan phylogeny. For example, to identify a diagnostic gene family with a signature duplication occurring just after the early metazoan radiation, we have searched for *Amphimedon* (sponge) or *Mnemiopsis* (ctenophore) genes with predicted 1-to-Many relationships with the other metazoan genomes. Remarkably, this strategy has not been productive, and at best has led to the identification of gene families with many possible root choices consistent with either a sponge-early or ctenophore-early model. A good example of this root intractability with an increasing number of deep gene duplications is the phylogeny of the metazoan WNT gene family (Fig. 1).

**Figure 1.**
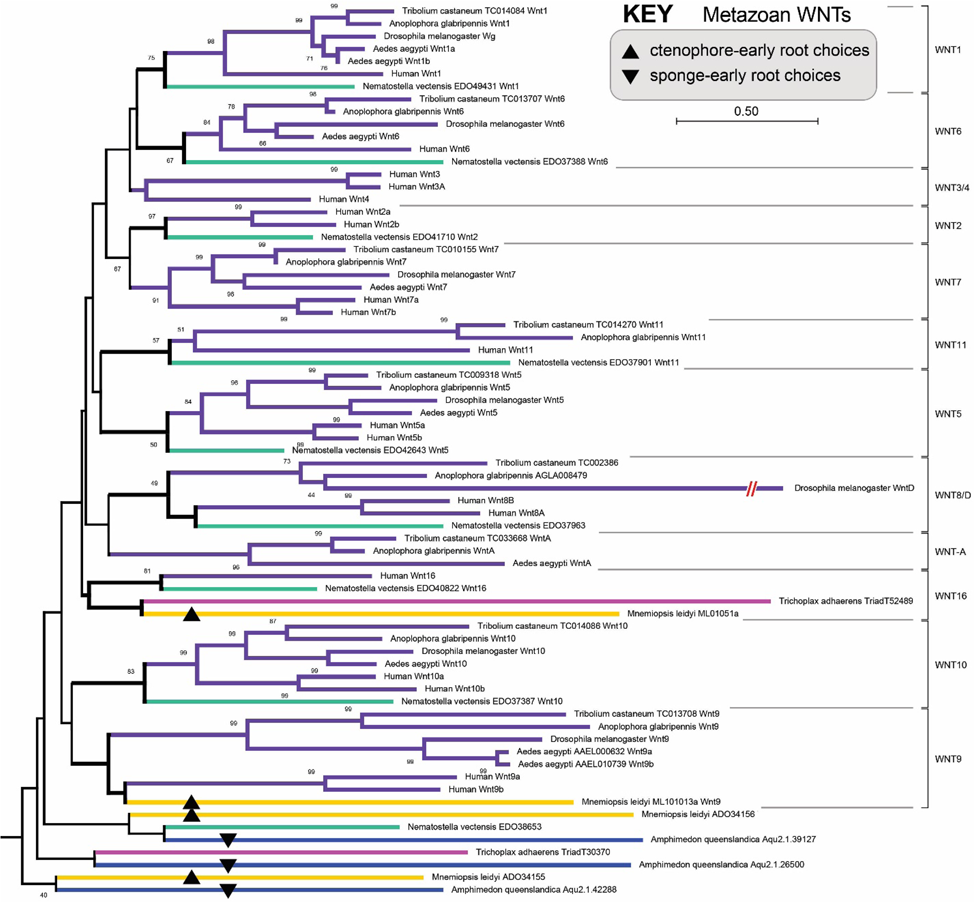
Early metazoan branching and the root of the WNT gene family. Example gene tree for the metazoan WNT ligand in which sponge and ctenophore genes exist in 1-to-Many relationship with other metazoan genes. A major problem in extracting a phylogenetic signal from a large gene family defined by many paralogs is the increasing ambiguity in identifying the true root of the tree, particularly when lacking a non-metazoan outgroup gene. For the WNT family tree shown here, there are four ctenophore-early (up arrowheads) and three sponge-early (down arrowheads) choices for rooting. In addition, there are other root choices for either model in combination with inferred gene losses. For example, here this tree is rooted between a weakly supported clade (40%) containing only sponge and ctenophore genes and the clade containing all the remaining genes. If this was the correct root, then there would have been a loss of a metazoan WNT paralog in the stem-eumetazoan lineage regardless of whether the true tree is a ctenophore-early or sponge-early tree. This tree was constructed using a distance-based method (Neighbor-Joining) with 100 boot strap replicate samples. Numbers indicate boot strap supports for values ≥ 40%. The same color scheme is used throughout this study as follows: violet = Bilateria, cyan = Cnidaria, magenta = Placozoa, blue = Porifera (sponges), and mustard/yellow/orange = Ctenophora (different hues for different species).

As a possible explanation for our failure to find unambiguous, early metazoan, gene paralogies, we hypothesized the existence of a methodological bias associated with the various ortholog-calling pipelines on which we were relying. This hypothesis of inherent bias in deciphering ancient gene paralogy motivated an alternative comparative genomic screen that we present here. We begin by illustrating the problems that can arise by not accounting for ancient paralogy.

We consider the consequences of a rooting procedure defined by a sponge-early model given an early-stem metazoan gene duplication event in either a true sponge-early world (Fig. 2A) or a true ctenophore-early world (Fig. 2B). These arguments are meant to illustrate that human and machine-predictions of orthology will encounter unique problems specific to the true animal sister lineage of all other animals particularly when phylogenies are constructed as “single copy” genes. (Many single-copy genes are actually members of much larger super-families and so are single-copy only in the sense that an analysis restricts itself to a sub-clade of genes).

**Figure 2.**
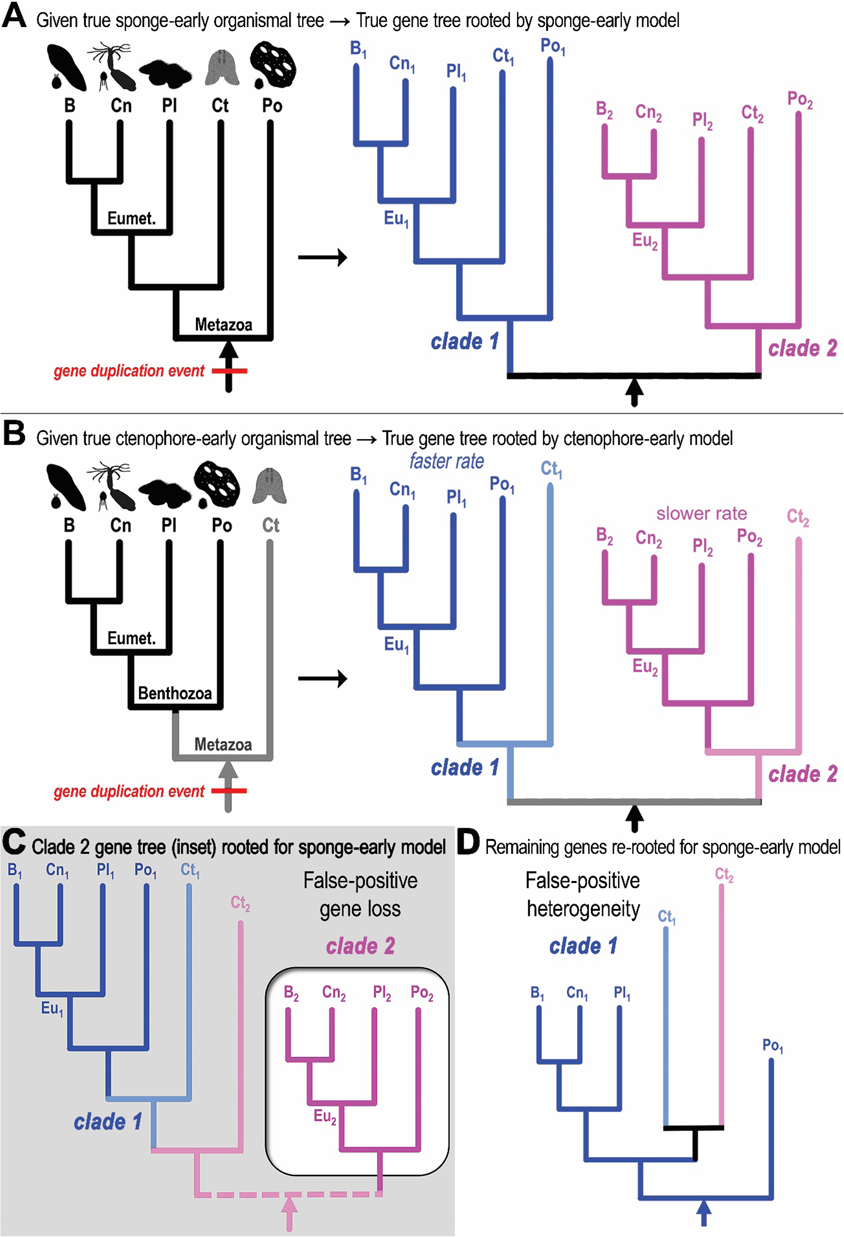
Dangers and promise of ancient paralogy for deducing the deep phylogeny of animals. **(A)** Given a true sponge-early phylogeny (left organismal tree with animal icons), a true gene tree for a gene duplicated in the stem-metazoan lineage (red bar) and maintained in each lineage without any gene loss is composed of two separate gene clades (blue clade one and magenta clade two), depicted here as evolving at different clade-specific rates. The principal metazoan lineages correspond to Bilateria (B), Cnidaria (Cn), Placozoa (Pl), Ctenophora (Ct), and Porifera (Po, sponges). Eumetazoa is defined here as Bilateria + Cnidaria + Placozoa. The true gene tree is rooted easily between the two clades and recapitulates the sponge-early model in each clade (Po_1_ and Po_2_ are the deepest branching lineages). **(B–D)** Depicted below the horizontal line are the consequences of forcing gene trees from a ctenophores-first world into a sponge-early model. **(B)** Shown is the competing ctenophore-early model in which Ctenophora is sister to all other animals (here referred to as Benthozoa as explained in the text). For the same stem-metazoan gene duplication depicted in A, we now have a corresponding true gene tree in which the ctenophore genes (Ct_1_ and Ct_2_) are the deepest branching lineages given the true ctenophore-early organismal tree. Asymmetric rates (faster or slower) are depicted in the gene phylogram, typical after divergent functionalization of paralogs. **(C)** The true gene tree in B is redrawn here with a different rooting procedure that isolates the more slowly-evolving paralog clade two (magenta clade in inset box) such that the sponge (Po_2_) lineage appears as the outgroup lineage within that subclade. This sponge-early re-rooting procedure maintains the topology of the true gene tree. The dotted line anticipates the analyses of the improperly partitioned subclades as separate “single-copy” gene clades. The partial clade two tree (inset) is missing the Ct_2_ ctenophore gene and would be associated with a false-positive gene loss signature. **(D)** The sponges-early model can then be applied to the remaining genes as shown. This tree includes the ctenophore Ct_2_ gene that was previously flipped into the faster evolving clade one, which also has a higher propensity for long-branch attraction than clade two. In contrast to the false-positive gene loss associated with the slowly-evolving clade two, the faster evolving clade one manifests false-positive heterogeneity of evolutionary rates and amino acid content due to the mixture of ctenophore paralogs.

In a sponge-early world, the sponge-early rooting procedure is simple and defines two gene clades (blue and magenta clades in Fig. 2A). The same is true if we use a ctenophore-early rooting procedure given a ctenophore-early world (Fig. 2B). We now describe the consequences of a forced sponge-early rooting given a ctenophore-early world, which has been the subject of much debate (Ryan *et al*. 2013; Moroz *et al*. 2014; Borowiec *et al*. 2015; Jekely *et al*. 2015; Whelan *et al*. 2015; Shen *et al*. 2017). For ease of reference, and for reasons explained in the Discussion, we will refer to the hypothetical sister clade composed of Eumetazoa + Porifera as “Benthozoa” (Fig. 2B).

By choosing the most closely-related homologs to the more conserved member of a pair of duplicated genes (the more slowly-evolving magenta subclade two of Fig. 2), or by internally re-rooting the true gene tree so that the subclade in question fits the standard model in which Porifera is sister to all other animals (Fig. 2C), we end up with an isolated subclade with a false-positive gene loss (inset box in Fig. 2C). Furthermore, the ctenophoran true outgroup sequence (“Ct_2_”) is easily collected into the more divergent sub-clade (blue subclade one in Fig. 2C). When the fast-evolving gene sub-clade is rooted separately with sponge as outgroup (“Po_1_”), the inherent topology unites both ctenophore paralogs in an apparent ctenophore-specific duplication (bottom phylogram in Fig. 2D). This second fast-evolving subclade would feature false-positive rate heterogeneity and compositional heterogeneity. False-positive compositional heterogeneity is consistent with recent attributions of (true) evolutionary sequence bias in ctenophores (Feuda *et al*. 2017).

As predicted by the ctenophore-early hypothesis (Fig. 2B) and deep paralogy-induced miscalling of orthologs (Fig. 2C–D), we find that relatively fewer metazoan orthologs are called in the ctenophore *Mnemiopsis leidyi* relative to sponge (*Amphimedon queenslandica*), placozoan (*Trichoplax adhaerens*), cnidarian (*Nematostella vectensis*), and bilaterian (the lophotrochozoan *Lottia gigantea* and *Capitella teleta*) genomes when orthologies are predicted according to a sponge-early model (Fig. 3). This finding supports the ctenophore-early hypothesis and is consistent with a similar approach in estimating deep animal phylogeny using gene content (Ryan *et al*. 2013).

**Figure 3.**
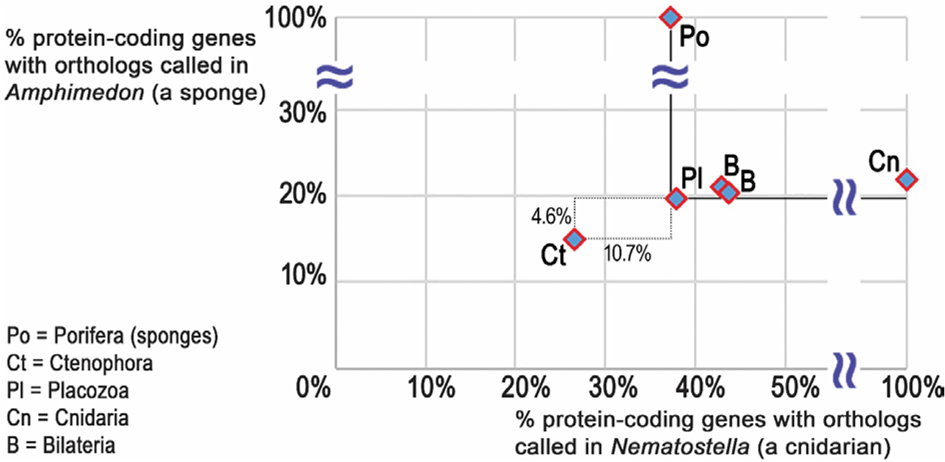
Deficit of metazoan homologs in ctenophores. Consistent with the predicted false-positive gene losses depicted in Fig. 2C, comparative genomic orthology data sets based on a sponge-early model result in far fewer orthologs called in a ctenophore relative to other metazoans. Shown are the percentages of each organism’s protein-coding gene set that have orthologs called in a cnidarian (*Nematostella*, *x*-axis) or a sponge (*Amphimedon*, *y*-axis) as inferred from the Metazoa ComparaEnsembl orthology calls (see Methods). Dotted box shows the distance from ctenophore value to the lowest value of the other animals along each axis (solid lines).

### A model-agnostic screen for early metazoan duplications

Based on the above observations and rationale, we devised a comparative genomic screen in which orthology calls to the ctenophore *Mnemiopsis* and the sponge *Amphimedon* are not considered during the initial candidate paralogy group screen. In other words, we relinquished our previous reliance on 1-to-Many predictions for sponge or ctenophore genes relative to other metazoans. We began with the 16,590 protein-coding genes from the beetle *Tribolium castaneum* (Tcas5.2 genes), which we chose as a model bilaterian that has not experienced extensive gene loss as in nematodes (Erives 2015), nor extensive gene duplications as in vertebrates. The *Tribolium* gene set becomes 8,733 genes if we only consider those that exist in paralogous relationship(s) with other *Tribolium* genes. Of these only 2,617 have orthologs called in the mosquito *Aedes aegypti*, the lophotrochozoan *Capitella teleta*, and the placozoan *Trichoplax adhaerens*. Then we discard those paralogous *Tribolium* genes which are related by a common ancestor in more recent taxonomic group such as Cucujiformia (an infraorder of beetles), Holometabola, Hexapoda, Mandibulata, Arthropoda, Pancrustacea, Protostomia, Bilateria, and Eumetazoa. We then grouped 2,038 candidate paralogous beetle genes into families of various sizes.

We considered the optimal size for a gene family to be used in resolving early animal branching order. First and foremost, having more duplications is likely to increase the probability that some of the duplications occurred during the early metazoan radiation that produced the extant animal phyla. However, choosing extremely large gene families containing many paralogous genes also introduces other problems in a classic trade-off scenario. For example, large gene families often present more intractable root choice problems, particularly when genes are absent in the outgroup lineage of choanoflagellates, as is the case for animal-only genes (*e.g.*, Fig. 1). Extremely large gene families seem to also have a greater rate of gene loss. This ‘*easily-duplicated, easily-lost*’ pattern further exacerbates the choice of root. We thus concluded that it is more effective to screen for gene paralogies defined by the smallest possible number of early metazoan gene duplications (two or three).

For the explained size trade-off reason, we set aside ∼56% (1,146 / 2038) of the beetle genes from the 153 largest paralogy groups, containing 4 to 41 genes each. We were then left with 892 genes grouped in < 382 small paralogy groups containing 2 to 3 genes each (Supporting Table S1). We sampled ∼16% (60 gene families) from these small candidate paralogy groups to find gene families that had at least one ortholog called in *Amphimedon queenslandica* and at least one in *Mnemiopsis leidyi*. We note that we cannot rule out the possibility that metazoan-relevant paralogies were excluded because true orthologs failed to be computationally called in sponges and ctenophores. Many of the candidate paralogy groups have paralogs called in sponges and ctenophores, indicating that the duplications occurred prior to the latest common ancestor (LCA) of Metazoa.

We constructed draft phylogenies using different approaches, including an explicit evolutionary model (Maximum Parsimony) and a distance-based method (Neighbor-Joining). If either of these initial trees indicated a possible early metazoan duplication, we then constructed trees using a more sophisticated and informative approach with metropolis-coupled-MCMC Bayesian phylogenetic inference in conjunction with more extensive gene curation of unannotated exons (see Methods). As detailed below, this new approach begins to identify choanozoan genes with early metazoan duplications. We present our first results for several gene families as demonstration of the value of our approach and in anticipation of finding more examples as additional genome assemblies from diverse metazoans become available.

### Duplication of the *cornichon*-family of transmembrane chaperones

As expected, many of the candidate gene families, which were identified without regard to presence or absence in *Amphimedon* or *Mnemiopsis*, were either: (*i*) missing in both ctenophores and sponges, or (*ii*) present in both ctenophores and sponges. The latter sometimes occurred when the ctenophore *Mnemiopsis* did not actually have the full set of duplications but a related ctenophore did. For example, in Figure 4A we show our initial draft tree for the Band 7/Stomatin and Stomatin-like paralogs. Initial trees suggested that this family was consistent with a signature of gene absence for Stomatin-like in the ctenophore *Mnemiopsis* (Order Lobata). However, subsequent follow-up shows that a ctenophore from a different order (Order Cydippida) has both paralogs (double asterisks). The clear presence of each paralog in at least one ctenophore means that this candidate could be set aside as being an uninformative duplication that occurred in an earlier choanozoan ancestor. The Stomatin/Stomatin-like example (Fig. 4A) also shows that our approach is likely to improve as additional genome assemblies become available for sponges and ctenophores. Nonetheless, other trees show from our screen, which was based on an agnostic preference for gene homologs in sponges and ctenophores, were in range to start being informative.

**Figure 4.**
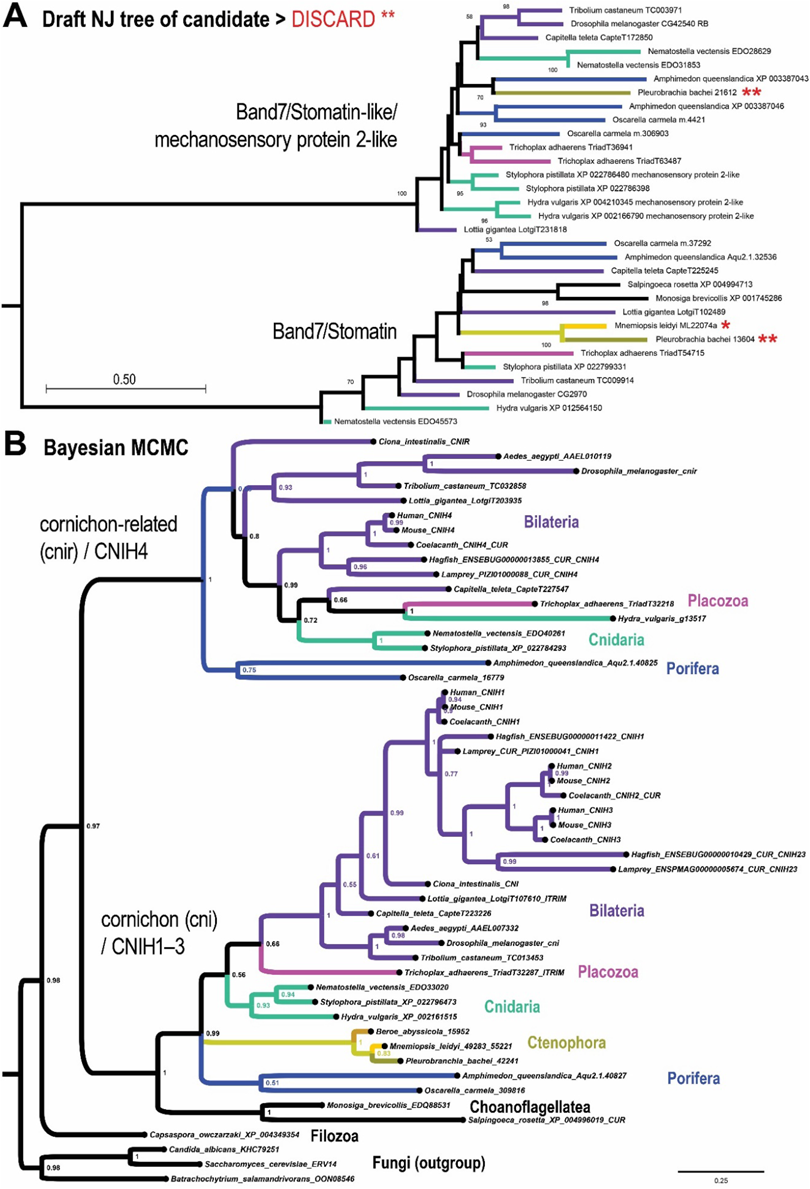
Example choanozoan duplications: *Stomatin* / *Stomatin-like*, and *cornichon* / *cornichon-related*. **(A)** Shown is the draft NJ tree for the Band 7/Stomatin and Stomatin-like paralogs, which was identified in our screen due to the signature of gene loss for Stomatin-like in the ctenophore *Mnemiopsis* (Order Lobata), which was indicated to have only a single Stomatin gene (single asterisk). Subsequent follow-up showed that a ctenophore from a different order (Order Cydippida) has both paralogs (double astrerisks). The clear presence of each paralog in at least one ctenophore means that this candidate can be set aside as being an uninformative duplication. Boot strap support is shown for values ≥ 40% from 500 replicates. **(B)** Shown is another example gene tree from our screen to identify candidate duplications based on a signature of gene loss (either true or false-positive gene loss). This gene family encodes a eukaryotic transmembrane receptor chaperone, which was either lost in both choanoflagellates and ctenophores (see *cnir* part of tree) as shown, or else was duplicated in the stem-benthozoan lineage and evolved sufficiently for it to be placed correctly in the tree. Tree was generated from a multiple sequence alignment composed of 167 alignment columns and is rooted with Holomycota (Fungi) as outgroup.

We find that the *cornichon* (*cni*/*CNIH1-CNIH3*) and *cornichon-related* (*cnir*/*CNIH4*) paralogy groups, which encode eukaryotic chaperones of transmembrane receptors (Herring *et al*. 2013), was either duplicated in the stem-choanozoan lineage with subsequent gene loss only in choanoflagellates and ctenophores, or else duplicated in the stem-benthozoan lineage with subsequent neofunctionalization. We find that both *cni* and *cnir* clades exist only in Porifera, Placozoa, Cnidaria, and Bilateria but not Ctenophora and Choanoflagellatea (Fig. 4B). This tree is based on a protein that is only ∼150 amino acids long and suggests that the ancestral *cni*/*cnir* gene was duplicated in the stem-choanozoan lineage with two subsequent independent losses in the stem-ctenophore lineage and the stem-choanoflagellate lineage (Fig. 4B).

The alternative interpretation of the *cni*/*cnir* duplication is that the progenitor gene was duplicated in the stem-benthozoic lineage with the *cnir* clade undergoing neofunctionalization to an extent promoting artifactual basal branching. In this interpretation there is no need to invoke independent gene losses for Ctenophora and Choanoflagellatea. Key differences in client proteins for the Cornichon chaperones (CNIH1/2/3) versus the Cornichon-related chaperone (CNIH4) support the neo-functionalization interpretation for the evolution of the CNIH4 paralog. Vertebrate CNIH1 and *Drosophila* Cornichon were found to mediate chaperone-like export of the transforming growth factor alpha (TGF-α)/Gurken precursor from the endoplasmic reticulum (ER) to the plasma membrane (Bokel *et al*. 2006; Castro *et al*. 2007), while the other vertebrate homologs of *Drosophila* Cornichon, CNIH2 and CNIH3, were found to mediate similar chaperone roles for AMPA receptor subunits (Schwenk *et al*. 2009). Both the TGF-α precursor and the individual AMPA receptor subunits have single transmembrane passes. In contrast, the vertebrate ortholog of Cornichon-related, CNIH4, has been found to mediate chaperone-like ER exit roles for certain G-protein coupled receptors (GPCRs), which possess seven transmembrane passes (Sauvageau *et al*. 2014). Thus, the Cornichon homologs, which are also present in choanoflagellates, may have evolved to help ER export of clients with only single transmembrane passes, while the Cornichon-related homolog CNIH4 may represent a neofunctionalization specialized for plasma membrane-bound proteins with multiple transmembrane passes.

In either case the progenitor *cni*/*cnir* gene was present as a single copy gene in the LCA of Holozoa as shown by the basal branching of the single gene from the filozoan *Capsaspora owczarzaki*, which branches before the duplication (Fig. 4B).

### TMC duplications define clades for Eumetazoa, Porifera, and Ctenophora

We find that a phylogenetic analysis of gene duplications of the transmembrane channels (TMCs), a large family of ion leak channels with roles in mechanotransduction (Delmas and Coste 2013; Pan *et al*. 2013; Ballesteros *et al*. 2018; Pan *et al*. 2018; Qiu and Muller 2018; Jia *et al*. 2019), excludes ctenophores from a eumetazoan super-clade composed of Bilateria, Cnidaria, and Placozoa (Fig. 5). We describe this gene family’s origin and diversification beginning with the deepest duplications in this family and then proceeding to the duplications most relevant to early metazoan evolution.

**Figure 5.**
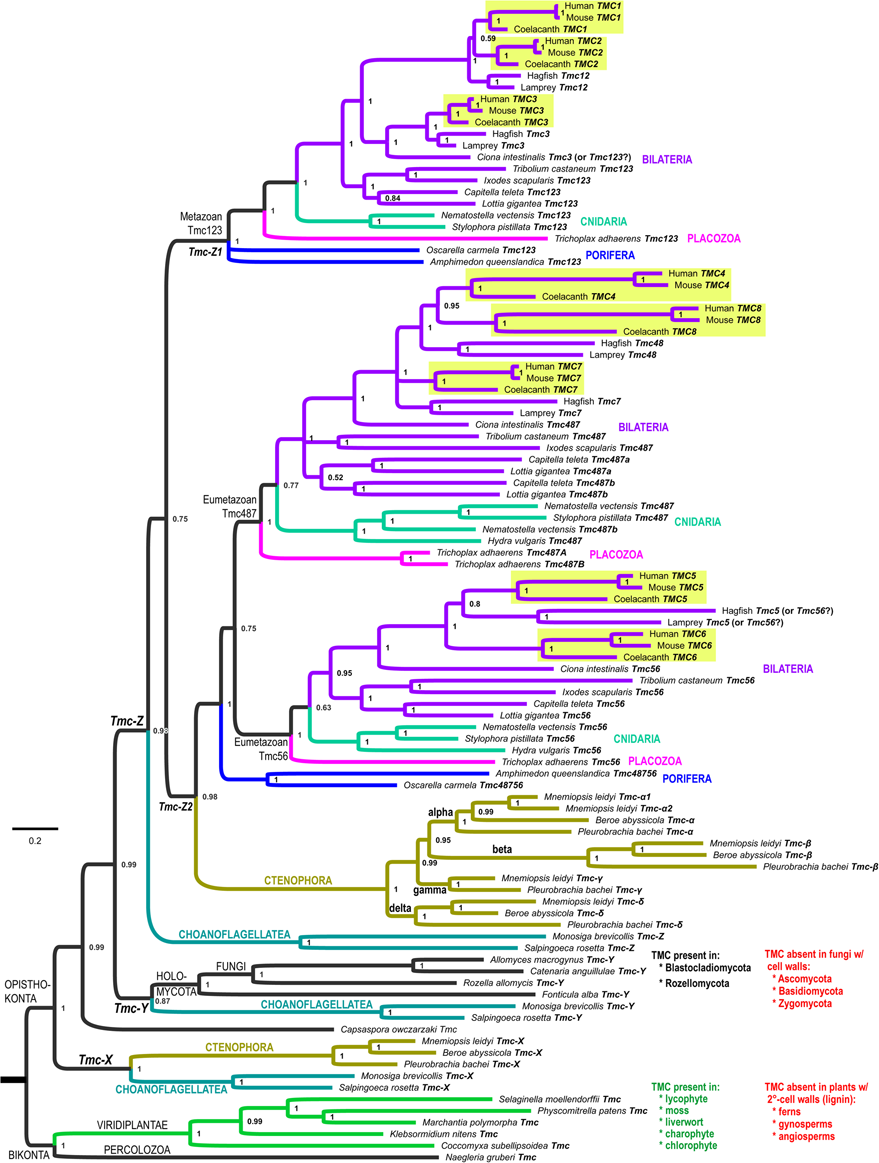
Independent duplications of *TMC* genes define separate clades for eumetazoans versus ctenophores. A phylogenetic tree of the mechanotransducing transmembrane channels (TMC) delineates early metazoan diversification as a branching of only three lineages: Eumetazoa (magenta Placozoa, teal Cnidaria, and violet Bilateria), Porifera (blue), and Ctenophora (gold shades). Each of the eight *TMC* orthogroups specific to gnathostomes are highlighted in yellow. This tree is based on a data set composed of 777 trimmed alignment columns and is rooted between unikonts (labeled “Opisthokonta” due to the apparent loss in Amoebozoa) and bikonts. Eumetazoan lineages share a duplication corresponding to the Tmc487 and Tmc56 TMC clades, while ctenophores share independent duplications of an ancestral Tmc48756 gene. The ctenophores share with choanoflagellates a more ancient Tmc duplication (labeled Tmc-X here).

In our TMC phylogeny, we identify genes from both Unikonta and Bikonta, which are the two main branches of the eukaryotic tree (Fig. 5). An affinity of the TMC proteins with OSCA/TMEM proteins has been proposed (Murthy *et al*. 2018) despite low sequence similarity (Ballesteros *et al*. 2018), but we find that the percent identity between predicted TMC proteins is higher than between TMC proteins and TMEM. For example, the gap-normalized percent amino acid identity in a pairwise alignment of human Tmc1 with the single *Naegleria gruberi* Tmc sequence is 17.3%, versus 12.7% (∼5% less) with human Ano3/TMEM16C (Supporting File S2). Moreover the identity between Tmc1 and TMEM16C goes away in a multiple sequence alignment suggesting it is inflated by the extra degree of freedoms allowed in a gapped pair-wise alignment.

In the context of this ancient eukaryotic provenance of the TMC family, we find an intriguing and suggestive pattern of TMC gene loss that is likely relevant to animal evolution. We find that the *Tmc* gene exists as a single copy in the genome of the non-amoebozoan slime mold *Fonticula alba* and a few early-branching fungal lineages, altogether representing the clade of Holomycota. One of these fungal lineages, *Rozella allomycis*, belongs to the Cryptomycota, a phylum notable for its absence of a chitinous cell wall typically present in most other fungi (Jones *et al*. 2011). The absence of a cell wall allows the Cryptomycota to maintain a phagotrophic lifestyle (Jones *et al*. 2011). The only other fungal lineages with a TMC gene are from Blastocladiomycota, suggesting that TMC genes were lost in all of Ascomycota, Basidiomycota, and Zygomycota, for which many genomes have been sequenced. Physiological incompatibility of Tmc function with certain cell walls is further supported by a similar distribution of Tmc genes in Viridiplantae (Fig. 5 green clade). We find that a single Tmc gene can be found in a chlorophyte (*Coccomyxa subellipsoidea*), a charophyte (*Klebsormidium nitens*), and in the multicellular plants of a liverwort (*Marchantia polymorpha*), a moss (*Physcomitrella patens*), and a lycophyte (*Selaginella moellendorfii*). The moss *Physcomitrella* and the lycophyte *Selaginella* represent a nonvascular land plant and a vascular land plant most closely-related to the clade composed of ferns, gymnosperms, and angiosperms, which correspond to the crown group having evolved lignin-based secondary cell walls (Sarkar *et al*. 2009). In addition, both the *Physcomitrella* and *Selaginella* genomes lack many of the cellulose synthase (*CesA*) genes responsible for the synthesis of plant cell wall components (Sorensen *et al*. 2010). Thus these TMC distributions in both plants and fungi suggest that rigid cell walls potentially interfere with physical environmental coupling to TMCs.

While we find TMC genes as a single copy in some lineages of the fungal and plant kingdoms, in *Naegleria gruberi*, and in the filozoan *Capsaspora owczarzaki*, we find that *Tmc* genes underwent duplication only during the holozoan radiation (or if earlier only with corresponding losses in fungi). In short, definitive TMC gene duplications can be found only within Choanozoa (choanoflagellates + Metazoa) (Fig. 5). The choanoflagellates *Monosiga brevicollis* and *Salpingoeca rosetta* have at least three TMC genes each, which we have labeled *Tmc-X*, *Tmc-Y*, and *Tmc-Z* for ease of reference here (Fig. 5). Remarkably, of all the animals, only ctenophores appear to have genes from the *Tmc-X* clade, which is a basally-branching Tmc clade that is sister to all of the remaining TMC genes from all of Opisthokonta. Only choanoflagellates and Holomycota have genes in the *Tmc-Y* clade. Last, except *Capsaspora*, whose single TMC gene is of uncertain affinity to the well supported *Tmc-X/Y/Z* clades, all of the remaining TMC genes are choanozoan genes from the *Tmc-Z* clade.

The TMC duplications within Choanozoa are informative for early choanozoan and metazoan branching. *Tmc-Z* apparently underwent a key duplication (“*Tmc<u>123</u>*” + “*Tmc<u>48756</u>*”) in the stem-metazoan lineage with a corresponding loss of one of the paralogs (“*Tmc<u>123</u>*”) in ctenophores. Alternatively, this duplication is actually a stem-benthozoan duplication that occurred after ctenophores split off from the rest of Metazoa. These two *Tmc-Z* subclades are so named here for their relationship to the vertebrate TMC1/2/3 (“*Tmc-Z1*”) and TMC4/8/7/5/6 (“*Tmc-Z2*”) paralogy groups. We find that this *Tmc-Z2* gene (*Tmc<u>48756</u>*) underwent a Eumetazoa-defining duplication that unites Placozoa, Cnidaria, and Bilateria (Fig. 5). Thus, in this stem-eumetazoan lineage we see that the *Tmc-Z2* bifurcates into the sister-clades of *Tmc<u>487</u>* and *Tmc<u>56</u>*. This duplication makes it highly improbable that Porifera or Ctenophora are more closely related to any single lineage within Eumetazoa. This contrasts with the stem-ctenophore lineage, in which the single *Tmc-Z2* gene independently duplicated several-fold to produce four ctenophore-specific duplications (*Tmc-α*, *Tmc-β*, *Tmc-γ*, and *Tmc-δ*) (Fig. 5, lineages in shades of yellow, gold, and mustard). This ctenophore specific repertoire is present in two different classes and three different orders of ctenophores. In Class Tentaculata, we have four *Tmc<u>48756</u>* genes from each of *Mnemiopsis leidyi* (Order Lobata) and *Pleurobrachia bachei* (Order Cydippida). In class Nuda, so named because of a complete (derived) loss of tentacles, we have only three *Tmc<u>48756</u>* genes from *Beroë abyssicola* (Order Beroida) because a representative gene from the *Tmc-γ* clade could not be identified.

Many metazoan sub-clades (*e.g*., Ctenophora, Lophotrochozoa, and Vertebrata) can be defined by clade-specific duplications. For example, lophotrochozoans share a duplication of *Tmc<u>487</u>* into *Tmc<u>487</u>a* + *Tmc<u>487</u>b*, while gnathostomes share a duplication of vertebrate *Tmc<u>12</u>* into *TMC1* + *TMC2* and *Tmc<u>48</u>* into *TMC4* + *TMC8*. If we include the cyclostomes (hagfish + lamprey), then all vertebrates share the duplication of *Tmc<u>123</u>* into *Tmc<u>12</u>* + *TMC3*, and *Tmc<u>487</u>* into *Tmc<u>48</u>* + *TMC7*. Thus, the “canonical” eight gene vertebrate repertoire represented by *TMC1*–*TMC8* is more correctly characterized as a gnathostome-specific *TMC* repertoire because cyclostomes are united in sharing only some of the duplications seen in humans (see Fig. 5).

Considering the *Tmc* gene duplications that are ancestral to Choanozoa and those specific to metazoan phyla, this phylogeny, based on the large TMC protein spanning ten transmembrane domains, unites Placozoa, Cnidaria, and Bilateria into a single Eumetazoa in the following ways. First, of greatest significance, is the shared derived eumetazoan duplication of *Tmc<u>48756</u>* into *Tmc<u>487</u>* and *Tmc<u>56</u>*. Second, is the close placement of Porifera as the sister group of Eumetazoa in both the *Tmc<u>123</u>* and *Tmc<u>48756</u>* subclades. Third is the unique situation that ctenophores possess TMC genes from an ancestral *Tmc-X* clade that were apparently lost in a stem-benthozoan lineage while also possessing an expansive repertoire via ctenophore-specific duplications (*Tmc-α*, *Tmc-β*, *Tmc-γ*, and *Tmc-δ*). These latter duplications are most parsimonious if they occurred in a metazoan lineage that branched off prior to the eumetazoan duplication within the Tmc-Z2 clade. In the Discussion, we speculate on the significance of the evolution of mechanotransduction in connection with the evolution of diverse ‘body plans’ during the metazoan radiation.

### A bHLH-ZIP duplication unites Benthozoa

We find that a new gene family from the bHLH-ZIP superfamily originated in the stem-metazoan lineage most likely from a duplication of the more distantly-related Max bHLH-ZIP gene (Fig. 6). This gene occurs as one gene copy in all ctenophores but as a pair of duplicated genes in Porifera and Eumetazoa (Fig. 6A). The pair of paralogous genes corresponds to *Max-like* X (*MLX*)/*bigmax* and *MLX Interacting Protein* (*MLXIP*)/*Mondo*. Mondo paralogs are also known as the carbohydrate response element binding protein (ChREBP) (Yamashita *et al*. 2001; Iizuka *et al*. 2004; Havula and Hietakangas 2018). The progenitor gene likely encoded a bHLH-ZIP homodimeric transcription factor (TF) in the stem-metazoan lineage, continuing as such into modern ctenophores, but evolving into the MLX:MLXIP heterodimer found in all other animals (Bigmax:Mondo in *Drosophila*, MLX:MondoA or MLX:MondoB in gnathostomes).

**Figure 6.**
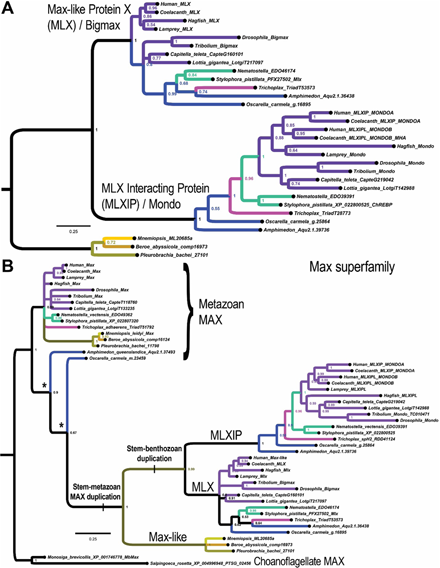
*MLX*/*bigmax* and *MLXIP*/*Mondo* encode a bHLH-ZIP heterodimer originating in a stem-benthozoan gene duplication. **(A)** Phylogenetic analysis of a MLX/MLXIP bHLH-ZIP gene family originating in the stem-metazoan lineage (possibly from a duplication of the distantly-related *Max* gene) followed by a second duplication after the divergence of the ctenophore lineage. Color coding of lineages follows Figure 3B except when topology precludes coloring sister stem lineages. This tree is rooted with ctenophores as the outgroup lineage. This tree is based on a data set composed of 263 alignment columns. **(B)** Shown is a phylogenetic tree constructed by Bayesian MC-MCMC and including sequences from the most closely-related and presumed progenitor MAX family (207 alignment columns). This tree places the Max-like sequences within the Max superfamily and furthermore puts the *MLX* and *MLXIP* duplications as occurring in a stem-benthozoan lineage that is sister to the Max-like sequence of choanoflagellates. While the *Max-like*/*MLX*/*MLXIP* sub-family likely originated as a stem-metazoan duplication, its sequences have diverged sufficiently to pull the sponge MAX sequences in a long-branch attraction (LBA) artifact (asterisks). This LBA is not the result of root choice because rooting between the choanozoan MAX sub-clade and the remaining sequences would result in sponges diverging prior to choanoflagellates, which would also correspond to LBA. Bayesian MCMC (3.2 M generations) finishing at ∼0.015 average standard deviation of split frequencies.

To identify the origin of the MLX and MLXIP bHLH-ZIP paralogs, we then conducted phylogenetic analyses with the most closely-related bHLH-ZIP sequences, which correspond to the MAX proteins. We find that the MLX family likely originated from an earlier stem-metazoan duplication of MAX that is absent in choanoflagellates and other holozoans (Fig. 6B). The Myc:Max gene regulatory network is known to have evolved in an early choanozoan stem-lineage (Brown *et al*. 2008; Young *et al*. 2011). When we root with choanoflagellate MAX as the out-group, we see that the MLX and MLXIP paralogs emerge from an earlier and presumably neofunctionalized duplication within the MAX family (see “stem-metazoan MAX duplication” in Fig. 6B). However, this duplication appears to have occurred after the lineage leading to modern ctenophores branched away from all other animals as ctenophores appear to have only the progenitor Max-like protein (see “stem-benthozoan duplication” in Fig. 6B).

We find that the bHLH domain from ctenophores shares residues with both subfamilies (pink vs. black background in Fig. 7). In the Discussion we speculate on a possible role for this gene duplication in a lifecycle synapomorphy for the proposed clade of Benthozoa.

**Figure 7.**
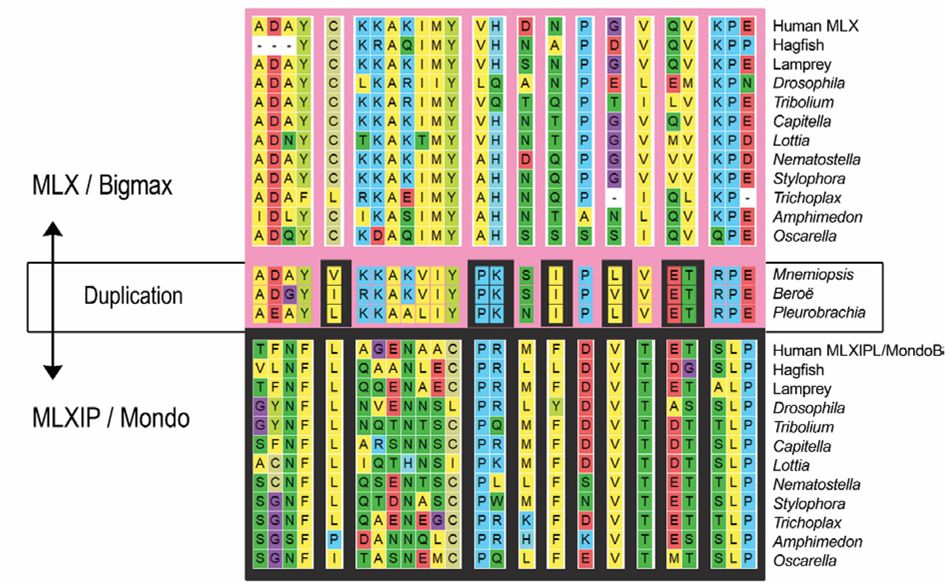
Shown are a subset of alignment columns in which the ctenophore residues more closely resemble either the MLX sequences (top pink) or the MLXIP/Mondo sequences (bottom dark gray). The MLXIPL/MondoB representative sequence is shown for humans.

## Discussion

By identifying and phylogenetically analyzing small paralogy groups that predate a eumetazoan super-clade composed of Bilateria, Cnidaria, and Placozoa, we find we can find gene duplications that are consistent with Ctenophora being the sister metazoan lineage to all other animals (Figs. 4–6). We discuss the significance of the different gene families that we identified and the possible relevance to metazoan biology.

We find that the transmembrane channel-like (TMC) gene family continued to duplicate and diversify for several phyla throughout metazoan evolution. For example, we see that the cyclostomes (hagfish and lampreys) branched early in the vertebrate tree prior to a number of duplications that occurred only in the stem-gnathostome lineage (*Tmc<u>12</u>* → *TMC1* + *TMC2*; *Tmc<u>48</u>* → *TMC4* + *TMC8*; *Mondo* → *MondoA*/*MLXIP* + *MondoB*/*MLXIPL*; and *CNIH<u>23</u>* → *CNIH2* + *CNIH3*). Nonetheless, cyclostomes and gnathostomes share the vertebrate-specific duplications corresponding to *Tmc12*, and *Tmc48* + *TMC7*. Thus, like the vertebrate clade and its subclades, many other metazoan phyla can be defined solely on the basis of *Tmc* gene duplications (Fig. 5). We thus propose that the evolutionary diversification of the Tmc channels throughout Metazoa, including the independent diversification within Ctenophora (Fig. 5), must have been under selection of two principal forces. The first is the changing physical constraints and sensorial opportunities associated with the evolutionary diversification of animal body plans themselves. The second factor is the evolutionary diversification of specialized cell types within their epithelial sensoria.

Mechanosensory, chemosensory and photosensory responses are universal among single and multicellular organisms and can be related to the evolution of specific proteins enabling already single cells to respond to mechanical, chemical and photic stimuli. For example, opsin proteins evolved in single-celled ancestors of metazoans (Feuda *et al*. 2012; Arendt 2017) and many single cell organisms can sense light, gravity and several chemical stimuli with dedicated sensors (Swafford and Oakley 2018). While the history of chemical and photic senses and the formation of specialized cell types and integration into appropriate organs in metazoans has been driven by the molecular insights into the molecular transducers (Arendt *et al*. 2016), mechanosensory transduction has seen less progress due to uncertainty of consistent association of a specific mechanotransducer channel across phyla (Beisel *et al*. 2010). On the one hand, mechanical sensation is clearly present in all single cell organisms to function as safety valves to release intracellular pressure sensed as tension in the lipid bilayer and was proposed as a possibly unifying principle of mechanosensation (Kung 2005). Follow up work showed a multitude of channels associated with mechanosensation (Beisel *et al*. 2010; Delmas and Coste 2013) arguing against a single unifying evolution of mechanotransduction.

Indeed, several families of mechanosensory channels have been identified whereby pores open as a function of lipid stretch or tethers attached to extra-or intracellular structures (Cox *et al*. 2018b; Cox *et al*. 2018a) Simply speaking, no single molecule has been identified to be associated with all mechanotransduction across phyla such as the ecdysozoan TRP channels also found in bony fish but absent in mammals (Beisel *et al*. 2010; Cox *et al*. 2018b; Qiu and Muller 2018).

Likewise, the ubiquitous Piezo mechanotransduction channels in Merkel cells (Delmas and Coste 2013; Ranade *et al*. 2015; Cox *et al*. 2016) were hypothesized to be the mechanotransduction channel of hair cells (Arendt *et al*. 2016) but have meanwhile been found not to be directly associated with the mechanotransduction process (Wu *et al*. 2017). Molecular analysis of mammalian mutations meanwhile has focused on the family of TMC (transmembrane channels) as possibly involved in hair cell mechanotransduction (Delmas and Coste 2013; Pan *et al*. 2013) and replacement of mutated TMC can restore hearing (Askew *et al*. 2015; Shibata *et al*. 2016; Yoshimura *et al*. 2018). More recently, molecular analysis has established that the TMC forms a mechanosensory pore as a homodimer with each subunit having ten predicted transmembrane domains (Pan *et al*. 2018). However, the detailed transmembrane protein dimer is unclear as other proteins are needed to transport the TMC channels to the tip (Pacentine and Nicolson 2019) and the entire complex of the vertebrate mechanosensory channel and its attachment to intra- and extracellular tethers remains unresolved (Qiu and Muller 2018). To what extent TMC family channel evolution aligns with mechanosensory cell and organ evolution remains to be seen but is already indicating unique features of ctenophores at every level (Fritzsch *et al*. 2006; Fritzsch *et al*. 2015). Recent work also establishes that ecdysozoan *Tmc<u>123</u>* paralogs function in body kinesthesia (proprioception), sensory control of locomotion or egg-laying behavior via membrane depolarization, and nociception (Guo *et al*. 2016; Wang *et al*. 2016; Yue *et al*. 2018). In summary, our study further lays the groundwork for understanding the molecular history of a sensory channel family in the context of the evolution of developmental gene regulatory circuits in different animal lineages (Corbo *et al*. 1997; Fritzsch *et al*. 2000; Beisel *et al*. 2010; Fritzsch and Elliott 2017; Cox *et al*. 2018a).

We speculate on the possible connection of the MLX + MLXIP/Mondo duplication (Fig. 6) to a transition from a holopelagic to a biphasic pelago-benthic life cycle *in the stem-benthozoan lineage*. This particular result furthers a growing picture of metabolic and chaperone gene loss and gene innovation in early animal evolution (Erives and Fassler 2015; Richter *et al*. 2018). Basic-helix-loop-helix (bHLH) TFs function as obligate dimers for DNA-binding (Murre *et al*. 1989; Lassar *et al*. 1991; Murre *et al*. 1991). Therefore, it is unlikely that a second MLX-related gene encoding a heterodimeric partner to the single gene found in ctenophores is artifactually missing in multiple sequenced genomes and transcriptomes (Ryan *et al*. 2013; Moroz *et al*. 2014). It is more likely that that the single *MLX-like* gene corresponds to the predicted evolutionary intermediate gene encoding a homodimeric bHLH TF.

The MLX:MLXIP/MLXIPL heterodimeric TF acts as a transcriptional regulator of metabolic pathways (*e.g.*, lipogenesis genes) in response to variation in intracellular sugar concentrations (Yamashita *et al*. 2001; Iizuka *et al*. 2004; Sans *et al*. 2006; Havula *et al*. 2013; Havula and Hietakangas 2018). In this regard it is interesting to speculate that the duplication evolved in connection with the evolution of a biphasic pelago-benthic life cycle featuring a pelagic feeding larva and a benthic feeding adult (whether it was a motile planula or sessile adult). Pelagic larval and benthic adult feeding forms would have distinct nutritional intakes associated with dissimilar feeding strategies and dissimilar nominal parameters. This bimodal variation would have evolved on top of variation associated with just a single feeding strategy. Thus, a stem-benthozoic ancestor may have demanded additional complexity in metabolic regulation that was subsequently afforded by duplicated paralogs to expand regulation of life cycle cell forms into differentiation of different cell types (Fritzsch *et al*. 2015).

### Methodological pitfalls of gene duplication as phylogenetic signal

While we may have identified some of the best candidates to date depicting early metazoan branching via gene duplications, we want to point out one possible pitfall in this approach. One of our candidates was the *Almondex* (*Amx*) superfamily of genes, which persisted in our screen through the stages where we required a missing orthology call for either the sponge *Amphimedon* or the ctenophore *Mnemiopsis* (see gene family #23 in size group 3 of Table S1). Almondex is a neurogenic TM2-domain containing protein characterized as a genetic modifier of Notch signaling in *Drosophila* (Shannon 1972; Shannon 1973; Michellod *et al*. 2003; Michellod and Randsholt 2008). Thus, as a neurogenic locus this candidate gene family could have made biological sense if it linked Ctenophora as a sister-group to Eumetazoa in the proposed Neuralia clade (Nielsen 2008). However, when we analyzed the Amx super-family in more depth, we found that one of the three paralogs was missing in the ctenophore *Mnemiopsis* (see CG11103 clade in Fig. 8), while all three paralogs were present in the sponge *Amphimedon* (Fig. 8).

**Figure 8.**
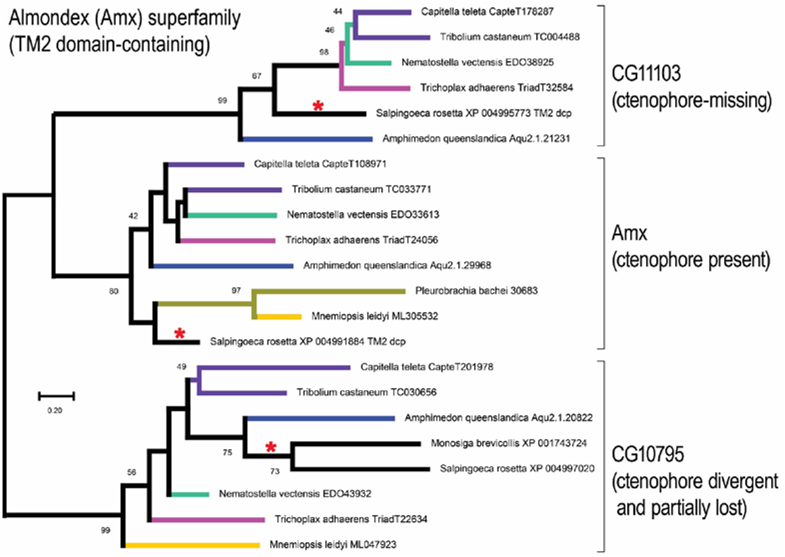
The neurogenic Almondex superfamily. Shown is a gene phylogeny composed of three paralogy groups (labeled by their *Drosophila* gene names), which originated in a choanozoan ancestor given the presence in both choanoflagellates (red asterisks) and animals. Given this ancient provenance, this tree may be rooted with any one of the three sub-clades as out-group. As explained in the text, ctenophores appear to be missing some of these paralogs (*e.g.*, CG11103), and were it not for the presence of each paralog in at least one choanoflagellate, alternate tree rooting strategies suggestive of gene duplications could have been possible. This tree was constructed using Neighbor-Joining (distance-based) with 1000 boot-strap replicates sampled from an alignment with 207 alignment columns. Boot-strap supports are shown only for values ≥ 40%.

To rule out *Mnemiopsis*-specific gene losses, we also searched for orthologs of each paralogy group in other ctenophores, and were indeed able to find Amx in the ctenophore *Pleurobrachia* (middle clade in Fig. 8). Superficially this tree began to resemble the MLX-MLXIP tree (Fig. 6B) except in its important relation to choanoflagellates. Unlike the single choanoflagellate *MAX* gene (Fig. 6B), the Amx superfamily tree shows that the choanoflagellate *Salpingoeca rosetta* has each of the three paralogs (red asterisks in Fig. 8). This result implies that ctenophores have lost the *CG11103* ortholog, and have either lost or are missing the *CG10795* ortholog (*Pleurobrachia*) or else possess a divergent version of this gene (*Mnemiopsis*).

In the case of the Amx superfamily, the evolutionary maintenance of Amx paralogs in at least one choanoflagellate is critical to ruling out duplicated genes, which would be possible with alternate root-choices. In the absence of the choanoflagellate *Salpingoeca* paralogs in the Amx and CG11103 sub-clades, which already appear to be lost in the choanoflagellate *Monosiga*, we may justifiably have rooted the tree in Figure 8 so that the choanoflagellate CG10795 clade is the outgroup. A tree rooted in this way (in the hypothetical absence of the two lone *Salpingoeca* sequences) allows the possibility that CG11103 could be a Benthozoa-specific duplication.

Thus, in using gene duplications to infer phylogenetic branching it is important to be mindful of unchecked misinterpretations facilitated by true gene loss. If this type of error is ubiquitous, that we should be able to find examples showing duplications shared by ctenophores and bilaterians even if they are false-positive duplications caused by unconstrained root choices. Further examples will be necessary to see if this is the case. But in summary, our comparative genomic screen designed to identify candidate families duplicated prior to the evolution of Eumetazoa has only identified examples consistent with a clade of ‘Benthozoa’ that unites Porifera and Eumetazoa as the sister-clade to Ctenophora.

## Author Contributions

A.E. and B.F. conceived this study together; A.E. conducted the computational analyses, including the phylogenomic tree constructions; A.E. and B.F. analyzed the data; A.E. wrote the manuscript, excluding the Discussion, which B.F. helped to write; and A.E. and B.F. edited the entire manuscript.

**Supplementary Information** is available in the online version of the paper.

## Supporting Information

**File S1.** Zipped file containing curation documents, fasta and alignment files for all gene sets.

**File S2.** TMC1/TMEM pairwise alignments.

**Table S1.** Candidate early metazoan paralogs. Shading in the first two columns alternates to highlight consecutive gene rows belonging to the same family.

